# Influence of GdVO_4_:Eu^3+^ nanocrystals on growth, germination, root cell viability and oxidative stress of wheat (*Triticum aestivum* L.) seedlings

**DOI:** 10.1101/2021.05.05.442722

**Authors:** Anna Ekner-Grzyb, Jagna Chmielowska-Bąk, Agata Szczeszak

## Abstract

Increasing application of lanthanide-doped nanocrystals (LDNCs) entails a risk of a harmful impact on the natural environment. Therefore, in the presented study the influence of gadolinium orthovanadates doped with Eu^3+^ nanocrystals on wheat (*Triticum aestivum* L.), chosen as a model plant species, was investigated. The seeds were grown in Petri dishes filled with colloids of LDNCs at the concentrations of: 0, 10, 50 and 100 µg/ml. The plants’ growth endpoints (number of roots, roots length, roots mass, hypocotyl length and hypocotyl mass) and germination rate were found to be not significantly changed after the exposure to GdVO_4_:Eu^3+^ nanocrystals at all used concentrations. The presence of LDNCs also had no effect on oxidative stress intensity determined on the basis of the amount of lipid peroxidation product (thiobarbituric acid reactive substances; TBARS) of the roots. Similarly, TTC (tetrazolium chloride) assay did not show any differences in cells’ viability. However, root cells of the treated seedlings contained less amount of Evans Blue (EB) when compared to the control.

## Introduction

Recently, a growing interest in the use of nanoparticles (NPs) has been observed. Their unique properties have been utilized in chemical and biological sciences, medicine, as well as in industry and agriculture. However, along with many advantages, their use entails some risks. They can enter the environment and may have impact on the wildlife and human health (1–3). The effects of NPs may be completely different than those of their larger counterparts (bulk materials) (4–7). These differences result from higher surface area to volume ratio and higher surface reactivity of nanostructures. Moreover, due to smaller size NPs could enter organisms more easily as compared to bulk structures ^4^, although it is not always the case ^5^.

NPs are taken up by plants and they are subsequently spread and accumulated in the tissues, mainly in roots, stems and leaves (8–11). The NPs influence the contaminated plants in many ways (12–14). They may negatively affect germination, cells morphology, growth rate or final biomass of plants and induce oxidative stress (6,15– 23). However, some authors have shown no negative consequences or even biostimulating effect of NPs on plants including oat (*Avena sativa* L.), corn (*Zea mays* L.), tomato (*Lycopersicon esculentum* L.), cress (*Lepidium sativum*), white mustard (*Sinapis alba*) sorghum (*Sorghum saccharatum*), lettuce (*Lactuca sativa* L.) and wheat (*Triticum aestivum* L.) (24–28). The positive effect occurred mainly at low concentrations and was reversed at higher doses of NPs (16,17). The improvement of growth may be caused by more effective uptake of water or nutrients in plants exposed to NPs (17,26,29), as well as interference with the toxic metals taken up from the medium (30,31). The overall effect of NPs on plants depends on numerous factors, such as time of the exposure, surface contamination, size, plants species, etc. (32– 35).

Inorganic nanocrystals doped with lanthanide ions (lanthanide-doped nanocrystals, LDNCs) have been widely investigated. Their unique spectroscopic properties, such as the ability to luminescence, resistance to photobleaching and photochemical degradation have been found very attractive for many potential applications (36,37). LDNCs are used in innovative technologies, optoelectronic industry (e.g. LCD), agriculture, as well as biological and medical sciences (e.g. drug delivery and bioimaging) (38–41). Among them, gadolinium-based LDNCs such as vanadates gadolinium doped with Eu^3+^ ions may be applied in magnetic resonance imaging, as catalysis, polarizers, laser hosts and luminescent materials (42).

Nanostructures used in the above-mentioned applications may be transferred to the environment. According to a study in California, the highest amount of gadolinium was observed in the water bodies close to hospitals and medical centres by the hospital and medical centres. This discovery suggests that the Gd-based contrast agents used for magnetic resonance imaging are the main source of contamination (43). Gadolinium may accumulate also in animal tissues, such as the human brain (44). A study with vertebrate cell lines has revealed that gadolinium may influence the vertebrate cells, e.g. cause apoptosis (45). Wysokińska et al. have reported that NaGdF_4_:Yb^3+^,Er^3+^, NaGdF_4_ or NaGdF_4_:Eu^3+^ NPs may negatively affect the viability of macrophages and fibroblast (39). Our previous study conducted on LDNCs similar to those used in this one has shown that Fe_3_O_4_@SiO_2_@GdVO_4_:Eu^3+^ 5% core@shell type nanostructure had no effect on the erythrocyte sedimentation rate, morphology of red blood cells or their membrane permeability when present in concentrations up to 1 mg/mL (46).

Not many authors have been interested in the toxicity of LDNCs towards plants. According to literature data, LDNCs may be taken up by particular parts of plants and their presence may influence, similarly like other nanostructures, plant development (9,10,16–18,47,48). Moreover, most of the previously published data on phytotoxicity of LDNCs concerned fluorides, mainly NaYF_4_ (9,10,16,17,49) or oxides (18). To the best of our knowledge, no data have been published on the phytotoxicity of vanadates gadolinium doped with europium ions. In the presented study the impact of GdVO_4_:Eu^3+^ NPs on wheat seedlings was investigated.

## Materials & Methods

### Nanoparticles characterisation and elemental analysis

The NPs used for ecotoxicity investigation were based on inorganic matrix of orthovanadates doped with Eu^3+^ ions, GdVO_4_: Eu^3+^, synthesized under hydrothermal conditions. Detailed description of the synthesis method and photophysical characterization of the NPs used are presented in the previous paper (50). After the synthesis of NPs, the concentration of the stock solution was 3.54 mg/ml as determined by inductively coupled plasma-optical emission spectrometry (Varian ICP-OES VISTA-MPX). Before the assays, transmission electron microscopy (TEM) images were recorded using an HRTEM JEOL ARM 200F transmission electron microscope at the accelerating voltage of 200 kV. Dynamic light scattering (DLS) and zeta potential measurements were performed using a Malvern Zetasizer Nano ZS instrument. Concentration of the colloid sample was 1 mg/mL. Zeta potential was measured at physiological pH.

In order to evaluate the uptake of LDNCs by the plant organisms, elementary analysis was used. After 3 days of incubation, the seedlings were collected, weighted, rinsed, and mineralized in a pressure reactor with HNO_3_. Next, the samples were analysed using inductively coupled plasma optical emission spectroscopy (ICP OES).

### Plant growth and treatment

Wheat (*Triticum aestivum* L.) seeds were kindly supplied by the Danko Company. The seeds were surface sterilised for 5 min. with 75% ethanol and another 10 min. with 1% sodium hyperchlorite according to (51). Afterwards, the seeds were washed with tap water for 30 min. and immersed in distilled water for 10 min.

Next, the seeds were incubated with the GdVO_4_:Eu^3+^ NPs according to (32) with some modifications. The seeds were placed in Petri dishes (30 seeds per dish) and filled with colloids of LDNCs at the following concentrations: 0, 10, 50 or 100 µg/ml of the aqueous solution. The concentrations were determined on the basis of the results of previous studies of lanthanide-doped nanostructures. The colloids were sonicated before all assays. They were grown in the dark at a stable temperature of 22°C for 3 days.

### Measurement of growth parameters

Germination rate, seedling growth (root and hypocotyl length), number of roots and fresh weight (root and hypocotyl mass) were measured. Six replicates (each consisting of 30 seedlings) were run per treatment.

### Cell viability estimation

Cell viability was evaluated by two spectrophotometric methods: staining with Evans Blue (EB) and tetrazolium chloride (TTC). The first method is based on the accumulation of EB in cells with damaged membrane. The measurements were performed according to the procedure given in (52) with some modifications. Approximately 200 mg of roots were cut off on the ice and incubated with 0.25% Evans blue (Sigma, E-2129) for 20 min. at room temperature. Afterwards, the roots were washed twice for 15 min. with distilled water, homogenized in a mortar with the distaining solution (50 ml of ethanol, 49 ml of distilled water, and 1 ml of 10% SDS) and incubated in the heating block for 15 min at 50°C. Next, the samples were centrifuged (12000 rpm, 20°C, 15 min.) and the absorbance of the supernatant was measured at 600 nm. Each experiment was repeated four times and all samples in each experiment were tested in duplicate.

The second method, consist in measurements of the amount of insoluble red formazan that is accumulated in living cells as a consequence of reduction of 2,3,5-triphenyltetrazolium chloride (TTC) with dehydrogenase (53). Seedling roots were cut off on ice and incubated with 1ml of 0.4% TTC (Serva; Heidelberg, Germany) for 24 hours at room temperature in the dark. Next, the roots were washed with distilled water, transferred to new tubes and flooded with 0.5 ml of 95% ethanol. The samples were homogenized with Tissue Lyser II (Qiagen) and incubated at 55°C for one hour. After that, another portion of 0.5 ml of 95% ethanol was added to the samples, which were next vortexed and centrifuged for 3 minutes at 10 000 rcf. The volumes of 800 µl of supernatant and 1400µl of 95% ethanol were mixed in spectrophotometer cuvette and the absorbance of the samples was measured at λ=485. Each experiment was repeated four times.

### Lipid peroxidation determination

For assessment of lipid peroxidation in seedling roots, the amount of thiobarbituric acid-reactive substances (TBARS) was measured, according to the method described in (54) with some modifications. Whole roots of seedlings (approximately 200 mg) were cut off on ice and homogenised with 3 ml of 10% TCA (Sigma-Aldrich, TO699). Thereafter, the samples were centrifuged (12000 rpm, 4°C, 10 min.) and 1 ml of supernatant was transferred to glass tubes, filled with 4 ml of 0.5% TBA dissolved in 10% TCA. After incubation at 95°C for 30 min., the samples were cooled. Next, they were mixed by inversion and centrifuged (5000 rpm, 4°C, 2 min.).

The absorbance of the supernatant was measured at 532 nm and corrected for unspecific absorbance at 600 nm (0.5% TBA in 10% TCA was used as a blank). Each experiment was repeated four times and all samples in each experiment were tested in duplicate.

### Statistics

Statistical analyses were performed using Statistica 13.3 for Windows (StatSoft). Differences in length and mass of the seedlings, as well germination rate between treatments were analysed with nonparametric Kruskall-Wallis test. For cell viability and lipid peroxidation quantification, analysis of variance (ANOVA) was used, followed by the Tukey post hoc test (if needed). Data were shown as a mean ± standard error (SE).

## Results

### Nanoparticles characterisation and elemental analysis

The NPs’ structure and morphology were characterized on the basis of DLS measurements and TEM images. DLS analysis showed that the hydrodynamic diameter of NPs, including their agglomerates, varies between 140 and 180 nm (Fig 1a). Furthermore, TEM image analysis confirmed that NPs form agglomerates with the average grain size 70 ± 5 nm. Additionally, a negative zeta potential (−20.0 ± 4,29 mV), indicates that NPs colloids are stable at physiological pH and appropriate to use as water colloids (55). To present stability and transparency of GdVO_4_:Eu^3+^ NPs aqueous colloids at concentration of 10 mg/ml, pictures in a daylight were taken (Fig 1b, left side). Furthermore, red luminescence under 254 nm UV excitation related to the presence of Eu^3+^ emitting ions was also performed (Fig 1b, right side).

**Fig 1.**
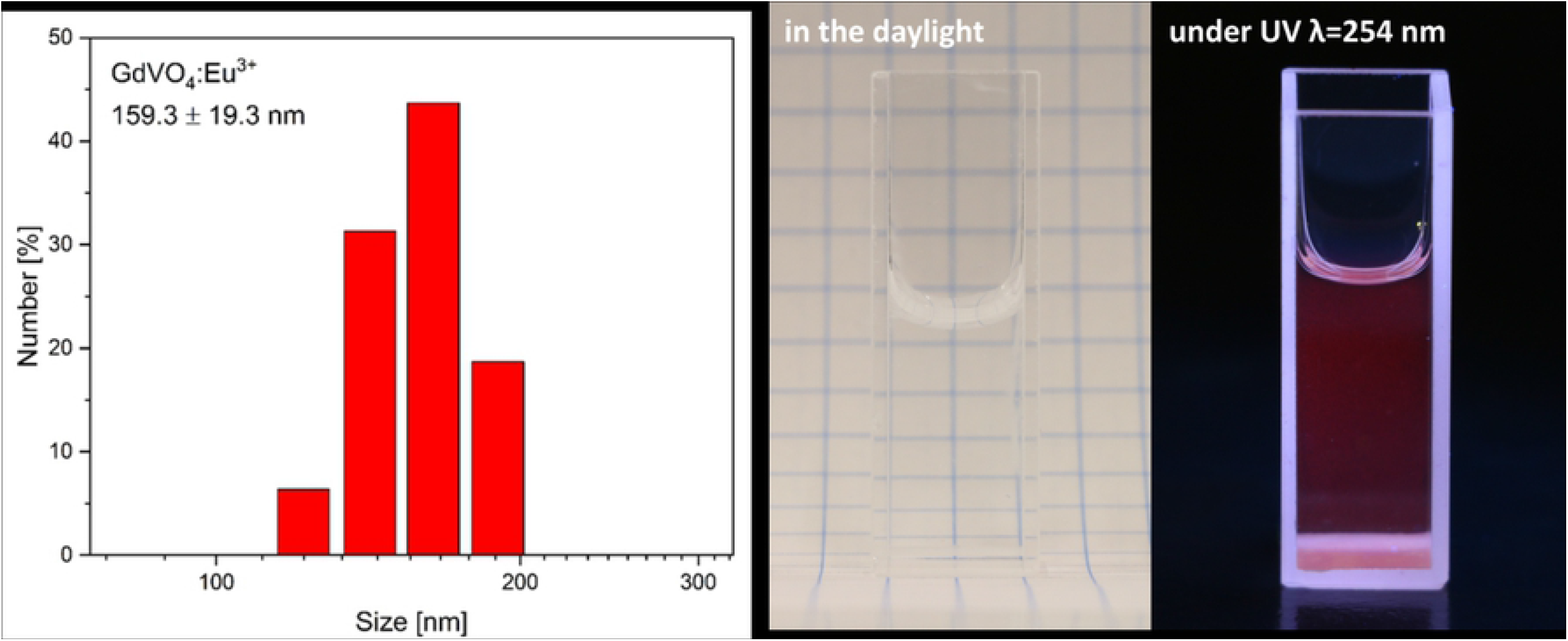
**a** DLS analysis of the GdVO_4_:Eu^3+^ NPs. **b** Photographs presenting the colloidal GdVO_4_:Eu^3+^ NPs, in a daylight (left) and under 254 nm UV light excitation (right)

ICP-OES analysis revealed that the LDNCs were taken up by the wheat seedlings.

### Growth and germination of the seedlings

Treatment with GdVO_4_:Eu^3+^ NPs had no significant impact on the morphology of wheat seedlings. The influence of the GdVO_4_:Eu^3+^ NPs on several growth endpoints, including the number of roots, roots length, hypocotyl length, roots mass and hypocotyl mass, was investigated. These parameters were not significantly changed as a result of the plants exposition for 3 days to 10, 50 and 100 µg/ml of LDNCs, relative to the controls (Kruskal-Wallis test, p>0.1; Fig 2a-e). The studied LDNCs also did not affect the germination rate of the tested concentrations (Kruskal-Wallis test, p>0.1; Fig 2f).

**Fig 2.**
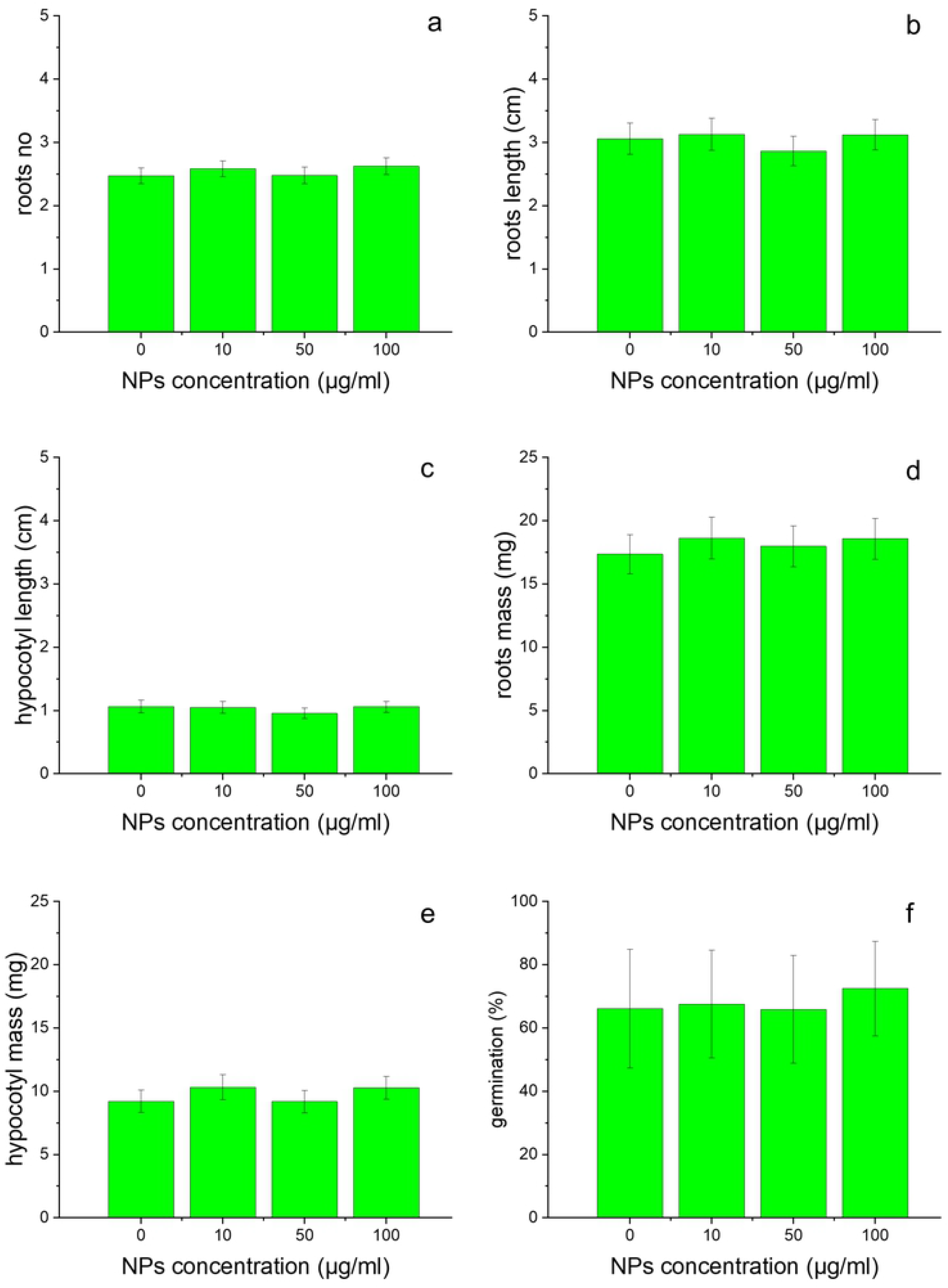
Number of roots (a), roots length (b), hypocotyl length (c), roots mass (d), hypocotyl mass (e) and germination rate (f) of wheat, treated for 3 days with the GdVO_4_:Eu^3+^ NPs at concentrations: 0, 10, 50 and 100 µg/ml. The results are presented as mean ± SE

### Oxidative stress

The degree of lipid peroxidation, estimated on the basis of the amount of TBARS did not change during the experiments. Treatments of the wheat with GdVO_4_:Eu^3+^ NPs did not affect this factor, at all tested concentrations (ANOVA, p=0.36; Fig 3).

**Fig 3.**
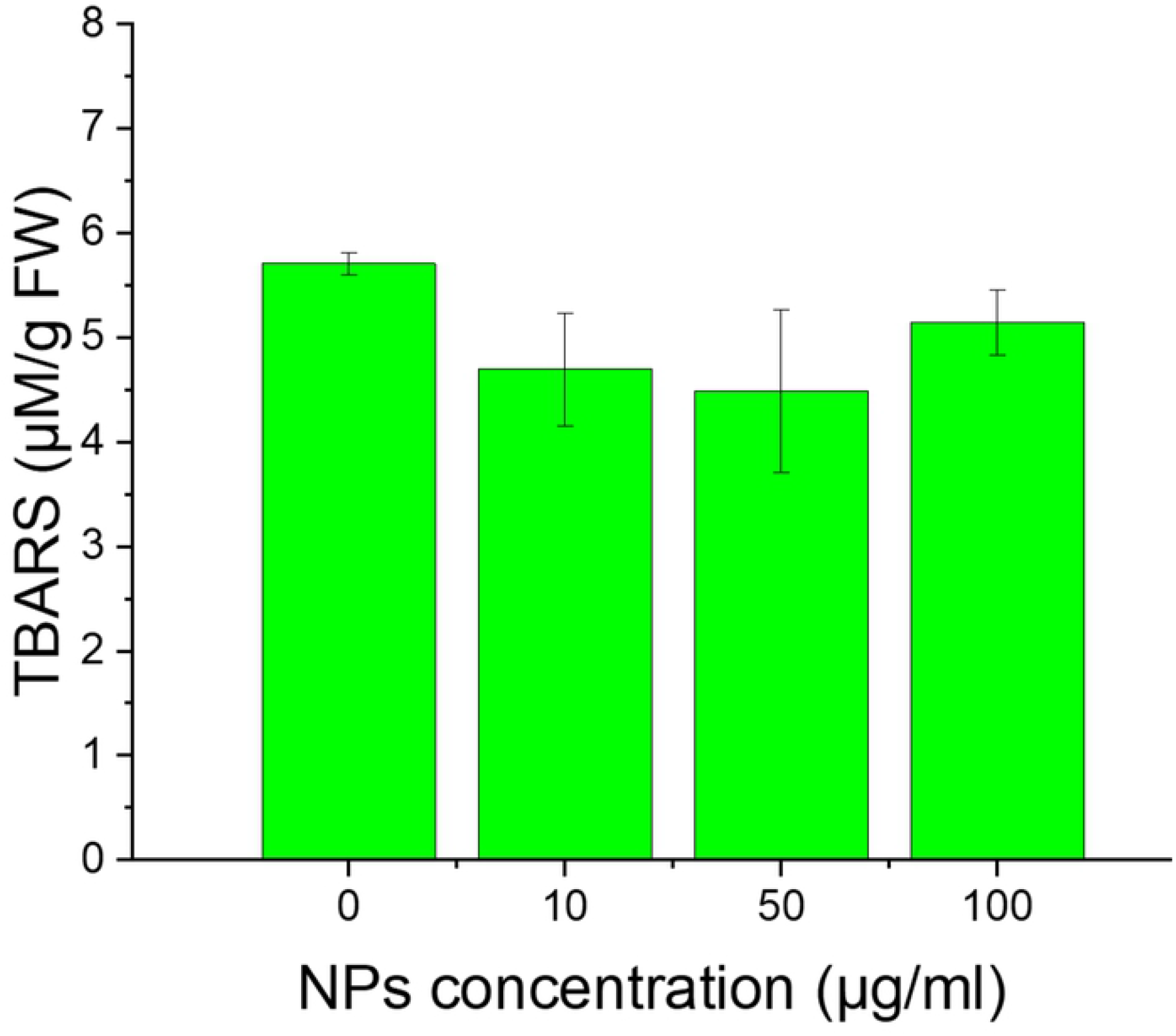
Lipid peroxidation of wheat cells, treated for 3 days with the GdVO_4_:Eu^3+^ NPs at concentrations: 0, 10, 50 and 100 µg/ml. The results are presented as mean ± SE

### Cell viability

The results of TTC staining showed that the exposure of the cells to GdVO_4_:Eu^3+^ NPs had no impact on the amount of formazan level (ANOVA, p=0.83; Fig. 4a). However, BE uptake was significantly different in control root cells and the cells incubated with LDNCs (ANOVA, p<0.005). The roots growing in the medium without LDNCs contained significantly more BE dye than those growing in the medium with NPs, both at the 10 µg/ml (Tukey test, p<0.05) and 50 or 100 µg/ml (Tukey test, p<0.01) concentrations (Fig 4b).

**Fig 4.**
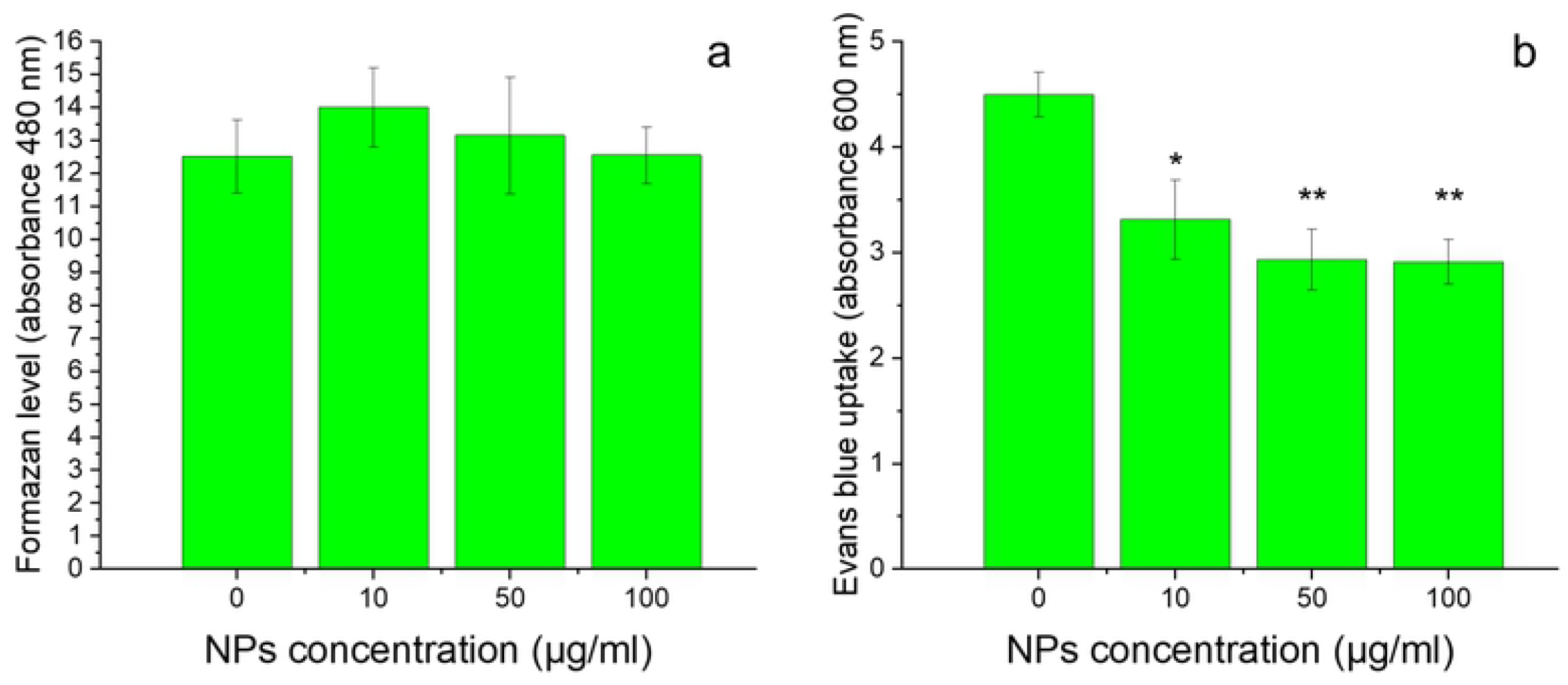
Cell viability of wheat exposed for 3 days to GdVO_4_:Eu^3+^ NPs at concentrations: 0, 10, 50 and 100 µg/ml, determined by TTC (a) and Blue Evans (b) staining assays. Asterisks (*) indicate statistically significant differences between the samples exposed to NPs and the control, Tukey test (*p<0.05 level, **p<0.01 level). The results are presented as mean ± SE

## Discussion

In the presented study, wheat seeds were incubated with GdVO_4_:Eu^3+^ NPs for 3 days. Even after this short time of treatment, the presence of LDNCs was detected inside the plants. However, results of the plants growth assays showed that GdVO_4_:Eu^3+^ NPs had no significant effect on germination rate, seedlings elongation and biomass of plants, when used in all tested concentrations (Fig. 2). To the best of our knowledge, there were no hitherto studies on the phytotoxicity of GdVO_4_: Eu^3+^ NPs. Although the LDNPs studied are poorly water soluble (56), some small amounts of the ions can be released to the solutions. Therefore, along with checking the influence of GdVO_4_:Eu^3+^ NPs on wheat plants we have to consider the effect of their particular compounds (e.g. ions). Similarly as we have found for wheat, Blinova et al. have suggested that the oxides containing gadolinium (Ce_0.9_Gd_0.1_O_2_, LaFeO_3_, Gd_0.97_CoO_3_) were not toxic for duckweed *Lemna minor* even at 100 µg/ml (57). According to other authors gadolinium caused growth inhibition of Chlorophyta (18,29), whereas Romero-Freire has suggested that this effect was observed only when Gd^3+^ ions were mixed with Ce^3+^ and Lu^3+^ lanthanide ions. In another study, Gd_2_O_3_ NPs caused a decrease in roots growth of wheat *Triticum aestivum* and rape *Brassica napus* (58). This effect depended on the sequence of processes of soaking and incubating during treatments. In the same study, germination of wheat and rape seeds was not affected by the presence of gadolinium oxides NPs (58). However, Gd^3+^ ions decreased significantly shoot growth of maize *Zea mays* at 10 µg/ml (59). Ruíz-Herrera et al. have shown that GdCl_3_ affected primary roots length in a dose-dependent manner (60). Unexpectedly, the same study has revealed that with increasing concentration of gadolinium ions, an increase in the total number of lateral roots per plant and lateral roots density increased (60). It has been reported that europium, another lanthanide component of the studied GdVO_4_:Eu^3+^ NPs, may be accumulated in macrophytes and alter the plant growth (61,62). The results obtained for one more component of the NPs studied, i.e. vanadium or vanadium oxide, which do not represent lanthanides, have been inconclusive. It has been suggested that plants incubated with high concentration of vanadium had shorter root length and poorer survival, whereas the low concentration of vanadium may improve the growth rate, accelerate flowering and biomass production (63), see the review in (64).

It has been demonstrated that modulated nutrient content in the tested plants may have impact on the growth rate. Garcia-Jimenez et al have reported that a solution of NH_4_VO_3_ added in a concentration of 5 mM stimulated the uptake of nutrients by the plants, when the concentration of this compound was increased to 15 mM, the stimulating effect was noticeable but reduced. Exposure to NH_4_VO_3_ at 5 µM has been found to cause an increase in the level of amino acids and sugars in leaves and roots (63). Other studies have revealed that the presence of europium caused a decrease in the concentration of macroelements in *Lathyrus sativus* roots (61). In the same paper it has been reported that europium activated proteolytic enzymes, and therefore increased the concentration of amino acids in the cells, causing plants to be less vulnerable to an arid environment (61). Saatz et al. have demonstrated significant correlations between the concentration of gadolinium in the nutrient solution and the concentration of Ca, Mg and P in the root tissues (59). However, it is unlikely that in the present study exposure to GdVO_4_:Eu^3+^ NPs could have impact on seedlings mineral composition, as the seedlings were grown in water (or colloids with LDNCs). The differences between literature data and our results on growth and development of wheat may be related to several factors, such as differences in solubility and structures of NPs, as well as assays’ duration. Moreover, the toxicity may depend on the presence of compounds other that the tested one (30,65).

The effect of NPs on lipid peroxidation (19,65–67), see the review in (13) and cell viability (8,32,66,68) has been studied, however, according to the present state of knowledge, the influence of LNDPs on the cells’ membrane damage and lipid peroxidation in plants has not been studied as yet. As mentioned above, the reported effects may be caused not only by LDNCs themselves but also by the ions released by these compounds. It has been shown that vanadium ions when present at the concentration 50 mg V in 1 kg of soil caused cell death, reduction of membrane integrity and activation of antioxidant enzymes in chickpea *Cicer arietinum* L. (69). In our study, the oxidative stress of the seedlings was evaluated by measuring the amount of lipid peroxidation product (thiobarbituric acid reactive substances, TBARS) as a biomarker. The assay showed that the GdVO_4_:Eu^3+^ NPs did not cause oxidation of the membrane lipid of the wheat root cells when used in all tested concentrations (Fig. 3).

The effect of the NPs presence on the cells viability was assessed by two methods: TTC and EB assays. They provided different results. According to the TTC results, the presence of LDNCs had no impact on the amount of formazan level, when used in all tested concentrations. It suggested that LDNCs did not cause cells death during the treatment. However, the results of the EB assay, revealed that incubation of the seedlings with GdVO_4_:Eu^3+^ NPs induced significant alterations in cells staining by EB dye. It has been proved that the presence of EB dye in cells is correlated with the loss of integrity of the plasma membrane of the cells and their mortality (52). Unexpectedly, in the presented study the control cells had higher absorbance than the treated ones (Fig. 4b). The mechanism of the observed phenomenon is not known, but some explanation can be proposed. First of all, the obtained results might be caused by decreased uptake of EB dye by the cells incubated with the LDNCs than the control. The LDNCs may create aggregates and therefore block pores of the cells, which prevents the access of the dye to the inside of the cells. It has been indeed shown that NPs might adhere to cell membranes and thus may alert intracellular transport (70). Alternatively, the LDNCs may interact with ions or structures responsible for pumping EB dye out.

## Conclusions

Summarising, there is no evidence for toxic effect of GdVO_4_:Eu^3+^ NPs on vascular plants, when used in concentrations of 100 µg/ml and lower. On the other hand, the observed decrease in the Evans Blue level in the treated seedlings indicated that GdVO_4_:Eu^3+^ NPs affect membrane functioning. Thus, it would be interesting to get more detailed insights into the effect of these NPs on plants physiology.

## Acknowledgements

This study was funded by Scientific Grant of the National Science Centre no. UMO-2016/23/D/NZ8/01112.

We are very grateful Danko Company for supplying wheat seeds.

## References

1. Judy JD, Bertsch PM. Bioavailability, Toxicity, and Fate of Manufactured Nanomaterials in Terrestrial Ecosystems [Internet]. 1st ed. Vol. 123, Advances in Agronomy. Elsevier Inc.; 2014. 1–64 p. Available from: http://dx.doi.org/10.1016/B978-0-12-420225-2.00001-7

2. Bystrzejewska-Piotrowska G, Golimowski J, Urban PL. Nanoparticles: Their potential toxicity, waste and environmental management. Waste Manag [Internet]. 2009;29(9):2587–95. Available from: http://dx.doi.org/10.1016/j.wasman.2009.04.001

3. Peralta-Videa JR, Zhao L, Lopez-Moreno ML, de la Rosa G, Hong J, Gardea-Torresdey JL. Nanomaterials and the environment: A review for the biennium 2008–2010. J Hazard Mater [Internet]. 2011 Feb;186(1):1–15. Available from: http://dx.doi.org/10.1016/j.jhazmat.2010.11.020

4. Hawthorne J, De La Torre Roche R, Xing B, Newman LA, Ma X, Majumdar S, et al. Particle-size dependent accumulation and trophic transfer of cerium oxide through a terrestrial food chain. Environ Sci Technol. 2014;48(22):13102–9.

5. De la Torre Roche R, Servin A, Hawthorne J, Xing B, Newman LA, Ma X, et al. Terrestrial Trophic Transfer of Bulk and Nanoparticle La 2 O 3 Does Not Depend on Particle Size. Environ Sci Technol [Internet]. 2015 Oct 6;49(19):11866–74. Available from: http://pubsdc3.acs.org/doi/10.1021/acs.est.5b02583

6. Modlitbová P, Novotný K, Pořízka P, Klus J, Lubal P, Zlámalová-GargoŠová H, et al. Comparative investigation of toxicity and bioaccumulation of Cd-based quantum dots and Cd salt in freshwater plant Lemna minor L. Ecotoxicol Environ Saf [Internet]. 2018 Jan;147(August 2017):334–41. Available from: http://linkinghub.elsevier.com/retrieve/pii/S0147651317305523

7. Lead JR, Batley GE, Alvarez PJJ, Croteau M-N, Handy RD, McLaughlin MJ, et al. Nanomaterials in the environment: Behavior, fate, bioavailability, and effects-An updated review. Environ Toxicol Chem. 2018 Aug;37(8):2029–63.

8. Ma Y, Zhang P, Zhang Z, He X, Li Y, Zhang J, et al. Origin of the different phytotoxicity and biotransformation of cerium and lanthanum oxide nanoparticles in cucumber. Nanotoxicology [Internet]. 2014;5390:1–9. Available from: http://www.ncbi.nlm.nih.gov/pubmed/24877678

9. Nordmann J, Buczka S, Voss B, Haase M, Mummenhoff K. In vivo analysis of the size-and time-dependent uptake of NaYF 4 :Yb,Er upconversion nanocrystals by pumpkin seedlings. J Mater Chem B [Internet]. 2015;3(1):144–50. Available from: http://xlink.rsc.org/?DOI=C4TB01515K

10. Hischemöller A, Nordmann J, Ptacek P, Mummenhoff K, Haase M. <I>In-Vivo</I> Imaging of the Uptake of Upconversion Nanoparticles by Plant Roots. J Biomed Nanotechnol [Internet]. 2009 Jun 1;5(3):278–84. Available from: http://openurl.ingenta.com/content/xref?genre=article&issn=1550-7033&volume=5&issue=3&spage=278

11. Soil C. Root System Architecture, Copper Uptake and Tissue Distribution in Soybean (Glycine max (L.) Merr.) Grown in Copper Oxide Nanoparticle (CuONP)-Amended Soil and Implications for Human Nutrition. Plants. 2020;9(1326):1–24.

12. Ma X, Geiser-Lee J, Deng Y, Kolmakov A. Interactions between engineered nanoparticles (ENPs) and plants: Phytotoxicity, uptake and accumulation. Sci Total Environ [Internet]. 2010 Jul 15;408(16):3053–61. Available from: http://dx.doi.org/10.1016/j.scitotenv.2010.03.031

13. Ma C, White JC, Dhankher OP, Xing B. Metal-Based Nanotoxicity and Detoxification Pathways in Higher Plants. Env Sci Technol. 2015;49(12):7109–22.

14. Kulacki KJ, Cardinale BJ. Effects of Nano-Titanium Dioxide on Freshwater Algal Population Dynamics. Bansal V, editor. PLoS One [Internet]. 2012 Oct 10;7(10):e47130. Available from: http://dx.plos.org/10.1371/journal.pone.0047130

15. Hao Y, Ma C, Zhang Z, Song Y, Cao W, Guo J, et al. Carbon nanomaterials alter plant physiology and soil bacterial community composition in a rice-soil-bacterial ecosystem. Environ Pollut [Internet]. 2018 Jan;232:123–36. Available from: https://doi.org/10.1016/j.envpol.2017.09.024

16. Yin W, Zhou L, Ma Y, Tian G, Zhao J, Yan L, et al. Phytotoxicity, Translocation, and Biotransformation of NaYF 4 Upconversion Nanoparticles in a Soybean Plant. Small [Internet]. 2015 Sep;11(36):4774–84. Available from: http://doi.wiley.com/10.1002/smll.201500701

17. Peng J, Sun Y, Liu Q, Yang Y, Zhou J, Feng W, et al. Upconversion nanoparticles dramatically promote plant growth without toxicity. Nano Res [Internet]. 2012 Nov 2;5(11):770–82. Available from: http://link.springer.com/10.1007/s12274-012-0261-y

18. Joonas E, Aruoja V, Olli K, Syvertsen-Wiig G, Vija H, Kahru A. Potency of (doped) rare earth oxide particles and their constituent metals to inhibit algal growth and induce direct toxic effects. Sci Total Environ [Internet]. 2017 Sep;593–594:478–86. Available from: http://dx.doi.org/10.1016/j.scitotenv.2017.03.184

19. Ma Y, Xie C, He X, Zhang B, Yang J, Sun M, et al. Effects of Ceria Nanoparticles and CeCl 3 on Plant Growth, Biological and Physiological Parameters, and Nutritional Value of Soil Grown Common Bean (Phaseolus vulgaris). Small. 2020;1907435:1–9.

20. Lee W-M, An Y-J, Yoon H, Kweon H-S. TOXICITY AND BIOAVAILABILITY OF COPPER NANOPARTICLES TO THE TERRESTRIAL PLANTS MUNG BEAN (PHASEOLUS RADIATUS) AND WHEAT (TRITICUM AESTIVUM): PLANT AGAR TEST FOR WATER-INSOLUBLE NANOPARTICLES. Environ Toxicol Chem [Internet]. 2008;27(9):1915. Available from: http://doi.wiley.com/10.1897/07-481.1

21. Oukarroum A, Barhoumi L, Pirastru L, Dewez D. Silver nanoparticle toxicity effect on growth and cellular viability of the aquatic plant Lemna gibba. Environ Toxicol Chem. 2013 Apr;32(4):902–7.

22. Andersen CP, King G, Plocher M, Storm M, Pokhrel LR, Johnson MG, et al. Germination and early plant development of ten plant species exposed to titanium dioxide and cerium oxide nanoparticles. Environ Toxicol Chem. 2016 Sep;35(9):2223–9.

23. Shu L, Yan Q, He Z, Xu K. Effects of Titanium Dioxide Nanoparticles on Photosynthetic and Antioxidative Processes of Scenedesmus obliquus. Plants. 2020;9(1748):1–13.

24. Libralato G, Costa Devoti A, Zanella M, Sabbioni E, Mičetić I, Manodori L, et al. Phytotoxicity of ionic, micro-and nano-sized iron in three plant species. Ecotoxicol Environ Saf [Internet]. 2016 Jan;123(2008):81–8. Available from: http://linkinghub.elsevier.com/retrieve/pii/S0147651315300312

25. Gavina A, Bouguerra S, Lopes I, Marques CR, Rasteiro MG, Antunes F, et al. Impact of organic nano-vesicles in soil: The case of sodium dodecyl sulphate/didodecyl dimethylammonium bromide. Sci Total Environ [Internet]. 2016 Mar;547:413–21. Available from: http://dx.doi.org/10.1016/j.scitotenv.2015.12.163

26. Khodakovskaya M, Dervishi E, Mahmood M, Xu Y, Li Z, Watanabe F, et al. Carbon Nanotubes Are Able To Penetrate Plant Seed Coat and Dramatically Affect Seed Germination and Plant Growth. ACS Nano [Internet]. 2009 Oct 27;3(10):3221–7. Available from: https://pubs.acs.org/doi/10.1021/nn900887m

27. Lian J, Wu J, Xiong H, Zeb A, Yang T, Su X, et al. Impact of polystyrene nanoplastics (PSNPs) on seed germination and seedling growth of wheat (Triticum aestivum L.). J Hazard Mater [Internet]. 2020 Mar;385:121620. Available from: https://doi.org/10.1016/j.jhazmat.2019.121620

28. Hayes KL, Mui J, Song B, Sani ES, Eisenman SW, Sheffield JB, et al. Effects, uptake, and translocation of aluminum oxide nanoparticles in lettuce: A comparison study to phytotoxic aluminum ions. Sci Total Environ [Internet]. 2020 Jun;719:137393. Available from: https://linkinghub.elsevier.com/retrieve/pii/S0048969720309037

29. Romero-Freire A, Joonas E, Muna M, Cossu-Leguille C, Vignati DAL, Giamberini L. Assessment of the toxic effects of mixtures of three lanthanides (Ce, Gd, Lu) to aquatic biota. Sci Total Environ [Internet]. 2019 Apr;661:276–84. Available from: https://doi.org/10.1016/j.scitotenv.2019.01.155

30. Mohmood I, Lopes CB, Lopes I, Tavares DS, Soares Amvm, Duarte AC, et al. Remediation of mercury contaminated saltwater with functionalized silica coated magnetite nanoparticles. Sci Total Environ [Internet]. 2016;557–558:712–21. Available from: http://www.sciencedirect.com/science/article/pii/S0048969716304995

31. Hu C, Liu L, Li X, Xu Y, Ge Z, Zhao Y. Effect of graphene oxide on copper stress in Lemna minor L.: evaluating growth, biochemical responses, and nutrient uptake. J Hazard Mater [Internet]. 2018 Jan;341:168–76. Available from: http://dx.doi.org/10.1016/j.jhazmat.2017.07.061

32. Silva S, Oliveira H, Craveiro SC, Calado AJ, Santos C. Pure anatase and rutile + anatase nanoparticles differently affect wheat seedlings. Chemosphere [Internet]. 2016 May;151:68–75. Available from: https://linkinghub.elsevier.com/retrieve/pii/S0045653516301990

33. Monteiro C, Daniel-da-Silva AL, Venâncio C, Soares SF, Soares Amvm, Trindade T, et al. Effects of long-term exposure to colloidal gold nanorods on freshwater microalgae. Sci Total Environ [Internet]. 2019 Sep;682:70–9. Available from: https://linkinghub.elsevier.com/retrieve/pii/S0048969719320637

34. Thiagarajan V M. P S. A R. S N. C G.K. S, et al. Diminishing bioavailability and toxicity of P25 TiO2 NPs during continuous exposure to marine algae Chlorella sp. Chemosphere [Internet]. 2019 Oct;233:363–72. Available from: https://doi.org/10.1016/j.chemosphere.2019.05.270

35. Silva S, Oliveira H, Silva AMS, Santos C. The cytotoxic targets of anatase or rutile + anatase nanoparticles depend on the plant species. Biol Plant [Internet]. 2017 Dec 3;61(4):717–25. Available from: http://link.springer.com/10.1007/s10535-017-0733-8

36. Mazierski P, Sowik J, Miodyńska M, Trykowski G, Mikołajczyk A, Klimczuk T, et al. Shape-controllable synthesis of GdVO 4 photocatalysts and their tunable properties in photocatalytic hydrogen generation. Dalt Trans. 2019;48(5):1662–71.

37. Wang D, Zhao C, Gao G, Xu L, Wang G, Zhu P. Multifunctional NaLnF4@MOF-ln nanocomposites with dual-mode luminescence for drug delivery and cell imaging. Nanomaterials. 2019;9(9).

38. Sun N, Ding Y, Tao Z, You H, Hua X, Wang M. Development of an upconversion fluorescence DNA probe for the detection of acetamiprid by magnetic nanoparticles separation. Food Chem [Internet]. 2018 Aug;257:289–94. Available from: http://linkinghub.elsevier.com/retrieve/pii/S0308814618303960

39. Wysokińska E, Cichos J, Kowalczyk A, Karbowiak M, Strzadała L, Bednarkiewicz A, et al. Toxicity mechanism of low doses of NaGdF 4 :Yb 3+, Er 3+ upconverting nanoparticles in activated macrophage cell lines. Biomolecules. 2019;9(1).

40. Dong R, Li Y, Li W, Zhang H, Liu Y, Ma L, et al. Recent developments in luminescent nanoparticles for plant imaging and photosynthesis. J Rare Earths [Internet]. 2019 Sep;37(9):903–15. Available from: https://linkinghub.elsevier.com/retrieve/pii/S100207211830930X

41. Szczeszak A, Grzyb T, Nowaczyk G, Ekner-Grzyb A. Emission colour changes in the CaF2 sub-microspheres doped with Yb3+, Er3+ and Mn2+ ions. J Alloys Compd [Internet]. 2020 Mar;817:152718. Available from: https://doi.org/10.1016/j.jallcom.2019.152718

42. Szczeszak A, Grzyb T, Śniadecki Z, Andrzejewska N, Lis S, Matczak M, et al. Structural, Spectroscopic, and Magnetic Properties of Eu 3+ -Doped GdVO 4 Nanocrystals Synthesized by a Hydrothermal Method. Inorg Chem. 2014 Dec;53(23):12243–52.

43. Hatje V, Bruland KW, Flegal AR. Increases in Anthropogenic Gadolinium Anomalies and Rare Earth Element Concentrations in San Francisco Bay over a 20 Year Record. Environ Sci Technol [Internet]. 2016 Apr 19;50(8):4159–68. Available from: https://pubs.acs.org/doi/10.1021/acs.est.5b04322

44. Kanda T, Fukusato T, Matsuda M, Toyoda K, Oba H, Kotoku J, et al. Gadolinium-based Contrast Agent Accumulates in the Brain Even in Subjects without Severe Renal Dysfunction: Evaluation of Autopsy Brain Specimens with Inductively Coupled Plasma Mass Spectroscopy. Radiology [Internet]. 2015 Jul;276(1):228–32. Available from: http://pubs.rsna.org/doi/10.1148/radiol.2015142690

45. Mizgerd JP, Molina RM, Stearns RC, Brain JD, Warner AE. Gadolinium induces macrophage apoptosis. J Leukoc Biol [Internet]. 1996 Feb;59(2):189–95. Available from: http://www.ncbi.nlm.nih.gov/pubmed/8603991

46. Szczeszak A, Ekner-Grzyb A, Runowski M, Mrówczyńska L, Grzyb T, Lis S. Synthesis, photophysical analysis, and in vitro cytotoxicity assessment of the multifunctional (magnetic and luminescent) core@shell nanomaterial based on lanthanide-doped orthovanadates. J Nanoparticle Res [Internet]. 2015 Mar 17;17(3):143. Available from: http://link.springer.com/10.1007/s11051-015-2950-4

47. Modlitbová P, Hlaváček A, Švestková T, Pořízka P, Šimoníková L, Novotný K, et al. The effects of photon-upconversion nanoparticles on the growth of radish and duckweed: Bioaccumulation, imaging, and spectroscopic studies. Chemosphere [Internet]. 2019 Jun;225:723–34. Available from: https://linkinghub.elsevier.com/retrieve/pii/S0045653519305107

48. Blinova I, Muna M, Heinlaan M, Lukjanova A, Kahru A. Potential Hazard of Lanthanides and Lanthanide-Based Nanoparticles to Aquatic Ecosystems: Data Gaps, Challenges and Future Research Needs Derived from Bibliometric Analysis. Nanomaterials [Internet]. 2020 Feb 14;10(2):328. Available from: https://www.mdpi.com/2079-4991/10/2/328

49. Hu S, Wu X, Chen Z, Hu P, Yan H, Tang Z, et al. Uniform NaLuF 4 nanoparticles with strong upconversion luminescence for background-free imaging of plant cells and ultralow power detecting of trace organic dyes. Mater Res Bull [Internet]. 2016 Jan;73:6–13. Available from: http://dx.doi.org/10.1016/j.materresbull.2015.08.020

50. Szczeszak A, Grzyb T, Śniadecki Z, Andrzejewska N, Lis S, Matczak M, et al. Structural, Spectroscopic, and Magnetic Properties of Eu 3+ -Doped GdVO 4 Nanocrystals Synthesized by a Hydrothermal Method. Inorg Chem [Internet]. 2014;53(23):12243–52. Available from: http://pubs.acs.org/doi/abs/10.1021/ic500354t

51. Chmielowska-Bąk J, Holubek R, Frontasyeva M, Zinicovscaia I, İşidoğru S, Deckert J. Tough Sprouting – Impact of Cadmium on Physiological State and Germination Rate of Soybean Seeds. Acta Soc Bot Pol [Internet]. 2020 Jun 24;89(2):1–10. Available from: https://pbsociety.org.pl/journals/index.php/asbp/article/view/asbp.8923

52. Jacyn Baker C, Mock NM. An improved method for monitoring cell death in cell suspension and leaf disc assays using evans blue. Plant Cell Tissue Organ Cult [Internet]. 1994 Oct;39(1):7–12. Available from: http://link.springer.com/10.1007/BF00037585

53. Towill LE, Mazur P. Studies on the reduction of 2,3,5-triphenyltetrazolium chloride as a viability assay for plant tissue cultures. Can J Bot [Internet]. 1975 Jun 1;53(11):1097–102. Available from: http://www.nrcresearchpress.com/doi/10.1139/b75-129

54. Heath RL, Packer L. Photoperoxidation in isolated chloroplasts. Arch Biochem Biophys [Internet]. 1968 Apr;125(1):189–98. Available from: https://linkinghub.elsevier.com/retrieve/pii/0003986168906541

55. Nuñez No, Rivera S, Alcantara D, de la Fuente JM, García-Sevillano J, Ocaña M. Surface modified Eu:GdVO4 nanocrystals for optical and MRI imaging. Dalton Trans [Internet]. 2013 Aug 14 [cited 2014 Apr 30];42(30):10725–34. Available from: http://www.ncbi.nlm.nih.gov/pubmed/23770723

56. Gnach A, Lipinski T, Bednarkiewicz A, Rybka J, Capobianco J a. Upconverting nanoparticles: assessing the toxicity. Chem Soc Rev [Internet]. 2015 Sep 1 [cited 2014 Oct 7];44(6):1561–84. Available from: http://www.ncbi.nlm.nih.gov/pubmed/25176037

57. Blinova I, Vija H, Lukjanova A, Muna M, Syvertsen-Wiig G, Kahru A. Assessment of the hazard of nine (doped) lanthanides-based ceramic oxides to four aquatic species. Sci Total Environ [Internet]. 2018 Jan;612:1171–6. Available from: https://doi.org/10.1016/j.scitotenv.2017.08.274

58. Ma Y, Kuang L, He X, Bai W, Ding Y, Zhang Z, et al. Effects of rare earth oxide nanoparticles on root elongation of plants. Chemosphere [Internet]. 2010 Jan;78(3):273–9. Available from: https://linkinghub.elsevier.com/retrieve/pii/S0045653509012764

59. Saatz J, Vetterlein D, Mattusch J, Otto M, Daus B. The influence of gadolinium and yttrium on biomass production and nutrient balance of maize plants. Environ Pollut [Internet]. 2015 Sep;204:32–8. Available from: https://linkinghub.elsevier.com/retrieve/pii/S0269749115001839

60. Ruíz-Herrera LF, Sánchez-Calderón L, Herrera-Estrella L, López-Bucio J. Rare earth elements lanthanum and gadolinium induce phosphate-deficiency responses in Arabidopsis thaliana seedlings. Plant Soil [Internet]. 2012 Apr 11;353(1–2):231–47. Available from: http://link.springer.com/10.1007/s11104-011-1026-1

61. Tian HE, Gao YS, Li FM, Zeng F. Effects of Europium Ions (Eu3+) on the Distribution and Related Biological Activities of Elements in Lathyrus sativus L. Roots. Biol Trace Elem Res [Internet]. 2003;93(1–3):257–70. Available from: http://link.springer.com/10.1385/BTER:93:1-3:257

62. Shtangeeva I. Europium and Cerium Accumulation in Wheat and Rye Seedlings. Water, Air, Soil Pollut [Internet]. 2014 Jun 27;225(6):1964. Available from: http://link.springer.com/10.1007/s11270-014-1964-3

63. García-Jiménez A, Trejo-Téllez LI, Guillén-Sánchez D, Gómez-Merino FC. Vanadium stimulates pepper plant growth and flowering, increases concentrations of amino acids, sugars and chlorophylls, and modifies nutrient concentrations. Niedz RP, editor. PLoS One [Internet]. 2018 Aug 9;13(8):e0201908. Available from: https://dx.plos.org/10.1371/journal.pone.0201908

64. Aihemaiti A, Gao Y, Meng Y, Chen X, Liu J, Xiang H, et al. Review of plant-vanadium physiological interactions, bioaccumulation, and bioremediation of vanadium-contaminated sites. Sci Total Environ [Internet]. 2020 Apr;712:135637. Available from: https://linkinghub.elsevier.com/retrieve/pii/S0048969719356323

65. Noman M, Shahid M, Ahmed T, Tahir M, Naqqash T, Muhammad S, et al. Green copper nanoparticles from a native Klebsiella pneumoniae strain alleviated oxidative stress impairment of wheat plants by reducing the chromium bioavailability and increasing the growth. Ecotoxicol Environ Saf. 2020;192(December 2019).

66. Silva S, Craveiro SC, Oliveira H, Calado AJ, Pinto RJB, Silva AMS, et al. Wheat chronic exposure to TiO2-nanoparticles: Cyto-and genotoxic approach. Plant Physiol Biochem [Internet]. 2017 Dec;121(October):89–98. Available from: https://linkinghub.elsevier.com/retrieve/pii/S0981942817303406

67. Yusefi-Tanha E, Fallah S, Rostamnejadi A, Pokhrel LR. Particle size and concentration dependent toxicity of copper oxide nanoparticles (CuONPs) on seed yield and antioxidant defense system in soil grown soybean (Glycine max cv. Kowsar). Sci Total Environ [Internet]. 2020 May;715:136994. Available from: https://doi.org/10.1016/j.scitotenv.2020.136994

68. Soares C, Branco-Neves S, de Sousa A, Pereira R, Fidalgo F. Ecotoxicological relevance of nano-NiO and acetaminophen to Hordeum vulgare L.: Combining standardized procedures and physiological endpoints. Chemosphere [Internet]. 2016;165:442–52. Available from: http://linkinghub.elsevier.com/retrieve/pii/S0045653516312437

69. Imtiaz M, Ashraf M, Rizwan MS, Nawaz MA, Rizwan M, Mehmood S, et al. Vanadium toxicity in chickpea (Cicer arietinum L.) grown in red soil: Effects on cell death, ROS and antioxidative systems. Ecotoxicol Environ Saf [Internet]. 2018 Aug;158:139–44. Available from: https://linkinghub.elsevier.com/retrieve/pii/S0147651318303154

70. Contini C, Schneemilch M, Gaisford S, Quirke N. Nanoparticle–membrane interactions. J Exp Nanosci [Internet]. 2018 Jan 1;13(1):62–81. Available from: https://doi.org/10.1080/17458080.2017.1413253

